# An analysis of semaphorin-mediated cellular interactions in the *C. elegans* epidermis using the IR-LEGO single-cell gene induction system

**DOI:** 10.1101/2024.01.24.577022

**Authors:** Motoshi Suzuki, Shin Takagi

## Abstract

One of the major functions of the semaphorin signaling system is the regulation of cell shape. In the nematode *C. elegans*, membrane-bound semaphorins SMP-1/2 (SMPs) regulate the morphology of epidermal cells via their receptor plexin, PLX-1. In the larval male tail of the SMPs/PLX-1 signaling mutants, the border between two epidermal cells, R1.p and R2.p, is displaced anteriorly, resulting in the anterior displacement of the anterior most ray, ray 1, in the adult male. To elucidate how the intercellular signaling mediated by SMPs regulates the position of the intercellular border, we performed mosaic gene expression analyses by using IR-LEGO (Infra<u>R</u>ed <u>L</u>aser <u>E</u>voked <u>G</u>ene <u>O</u>perator). We show that PLX-1 expressed in R1.p and SMP-1 expressed in R2.p is required for proper positioning of ray 1. The result suggests that SMPs signaling promotes extension, rather than retraction, of R1.p. This is in contrast to a previous finding that SMPs mediate inhibition of cell-extension of vulval precursor cells, another group of epidermal cells of *C. elegans,* indicating the context-dependence of cell shape control via the semaphorin signaling system.

## INTRODUCTION

Semaphorin is a major group of signaling molecules of the animal conserved both in vertebrates and invertebrates (Jongbloets and Pasterkamp, 2014). Semaphorins and the receptor plexins are involved in regulation of diverse developmental and physiological processes including neural circuit formation, vascular morphogenesis, bone development, tumorigenesis and regulation of immuno-response (Tran et al., 2007; Pasterkamp, 2012; Iragavarapu-Charyulu et al., 2020; Kang and Kumanogoh, 2013; Tamagnone, 2012; Kumanogoh and Kikutani, 2013). In axon guidance, semaphorin was first identified as a repellent for the growth cone, but later studies showed that the response of the growth cone to semaphorin is context-dependent and may turn to attraction depending on factors internal and external to the cell, such as second messengers, the FAK/src signaling, CNG channel or proteoglycans (Song et al., 1998; Falk et al., 2005; Togashi et al., 2008; Kantor et al., 2004).

A nematode *Caenorhabditis elegans (C. elegans)* has three semaphorins and two plexins. In larvae, Class I transmembrane-type semaphorins (SMP-1 and SMP-2: SMPs) bind to PlexinA (PLX-1) to regulate the shape of epidermal cells that generate adult organs such as the vulva in hermaphrodites and rays, sensory structures, in the male tail (Fujii et al., 2002; Ginzburg et al.,2002; Dalpé et al., 2004, 2005; Liu et al., 2005). Defective SMPs signaling affects the arrangement of larval epidermal cells. In vulval precursor cells (VPCs) of the SMPs-PLX-1 signaling mutants, VPCs fail to terminate the longitudinal extension, resulting in the overshoot and lateral overlaps among VPCs (Liu et al., 2005). In the larval male tail, the defective SMPs signaling, mainly mediated by SMP-1, causes an anterior shift in the position of the border between a pair of epidermal cells called R1.p and R2.p, which eventually results in anterior displacement of the anterior-most ray, ray 1, in the adult (Fujii et al., 2002).

While the SMPs-PLX-1 system appears to mediate a stop signal for cell extension in VPCs (Liu et al., 2005; Suzuki et al., 2022), little is known about how the semaphorin-plexin signal is exchanged among epidermal cells of the larval male tail and regulates their behavior. The receptor PLX-1 is expressed in all epidermal ray precursor cells, R(n)s, and their descendants including R1.p and R2.p (Fujii et al., 2002). Thus, although it is plausible that SMPs signaling between R1.p and R2.p keeps the border between them at a proper position, it is not clear how and in which of these two adjacent cells the functions of SMPs and PLX-1 molecules are required: the anterior shift of the border between R1.p and R2.p observed in the SMPs-PLX-1 signaling mutants can be explained either by the failure of retraction of the anterior part of the posterior cell, R2.p, or the failure of extension of the posterior part of the anterior cell, R1.p. While in these simple models SMPs, transmembrane-type semaphorins, and PLX-1 are assumed to interact in a cell-to-cell contact-mediated manner, this has not been tested experimentally; a previous study suggested that SMPs expressed in the hook primordium, which is located on the ventral midline distant from R1.p and R2.p, determine the position of ray 1 (Dalpé et al., 2004).

In order to address this issue, it would be useful to induce expression of PLX-1 and/or SMPs, SMP-1 in particular, in a single R(n) or its single descendant cell of each mutant, and examine the ray 1 phenotype. Whereas tissue-specific promoters, such as the promoter of *mab-5*, *egl-5*, *lin-32* and *efn-4* (Salser and Kenyon, 1996; Ferreira et al., 1999; Portman and Emmons, 2000; Hahn and Emmons, 2003) could be used to force expression of the transgenes in R(n)s and their descendant cells, the expression is not restricted to a single cell and the timing of gene expression cannot be regulated as we desire. To express PLX-1 and SMP-1 in a targeted single cell at a desired developmental stage, here we used IR-LEGO (<u>I</u>nfra<u>R</u>ed <u>L</u>aser <u>E</u>voked <u>G</u>ene <u>O</u>perator) (Kamei et al., 2009; Suzuki et al., 2013). Our mosaic gene expression analysis indicates that expression of PLX-1 in R1.p and expression of SMP-1 in R2.a or R2.p at a certain period of development is important for proper positioning of the border between R1.p and R2.p, suggesting that the SMPs signal promotes the extension of a PLX-1-expressing R1.p.

## MATERIALS AND METHODS

### C. elegans strains

Standard techniques for *C. elegans* culture and genetics were used as described by Brenner (1974).

The following strains were used.

N2, [LGI] *smp-1(ev715)*, *smp-2(ev709)*; [LGIV] *plx-1(nc37)*, *him-8(e1489)* [LGV] *him-5(e1490)*.

The transgenes used in this study include

jcIs1[ajm-1::gfp; rol-6](su1006)] (Mohler et al., 1998),
ncIs13[ajm-1::gfp] (Liu et al., 2005),
ncIs17[hsp::gfp] (Kamei et al., 2009),
ncIs19[hsp::plx-1; hsp::gfp; rol-6(su1006)] (Suzuki et al., 2022),
ncIs23[hsp::smp-1; hsp::gfp; rol-6(su1006)] (Suzuki et al., 2022),
ncEx2030[lin-32p::plx-1; rol-6(su1006)] (Nukazuka et al, 2008).
ncIs24[hsp::smp-1; hsp::gfp; rol-6(su1006)] was generated similarly to ncIs23 (Suzuki et al., 2022).
ncIs211[hsp::gfp] was generated similarly to ncIs17 (Kamei et al., 2009).
ncIs210[hsp::plx-1;hsp::smp-1; hsp::gfp; rol-6(su1006)] is a chromosomally integrated

transgene generated by the gamma-ray irradiation (Shioi et al., 2001) to *ncEx2016[hsp::plx-1; hsp::smp-1; hsp::gfp; rol-6(su1006)]* worms (Suzuki et al., 2022).

To generate the rescue construct *lin-32p::smp-1,* a *smp-1 cDNA* was cloned in the *pPD49.26* vector (provided by A. Fire) carrying the *lin-32* promoter *(lin-32p)* (Nakao et al., 2007). The plasmid was mixed with *pRF4* in the ratio of 1:2 (in weight) and was used to generate *ncEx2038[lin-32p::smp-1; rol-6(su1006)]* by microinjection (Mello and Fire, 1995).

For analyses of male tails, the strains carried the *him-5* or *him-8* mutation. The strains used for the IR-irradiation experiments contained *ajm-1::gfp* for visualizing the ray-lineage cells. Some stains also carried *rol-6(su1006),* which confers the Roller phenotype, as a transgene to facilitate the observation of rays.

NW1358 *smp-1(ev715)* and NW1335 *smp-2(ev709)* were provided by Joe Culotti (Ginzburg et al., 2002) and SU93 *jcIs1[ajm-1::gfp; rol-6(su1006)]* were provided by Jeff Simske (Mohler et al., 1998).

### IR-laser irradiation

Live males at the mid-L3 stage were placed on a 6% agar pad mounted with 5% levamisole for immobilization, and covered with a coverslip with a thickness of approximately 0.2 mm (Matsunami, Japan). The IR-LEGO system (Olympus IX-70 equipped with the IR-laser set) was described in the previous papers (Kamei et al., 2008; Suzuki et al., 2022). Specimens were observed with a 100 X mono-layer coated objective lens (UplanApo 100 X NA 1.3, Olympus, Japan) immersed in oil (Immersion oil type-F, Olympus, Japan). After irradiating worms, the coverslips were immediately removed and worms were recovered on a NGM plate. Three hours after irradiation, we checked GFP expression in the targeted cell and then observed rays one day after irradiation. In typical experiments, ray-lineage cells at stage 2 were irradiated with IR-laser beam at 13 mW for 0.25 sec a total of 4 times. About 15% of irradiated R(n).as and R(n).ps were induced to express GFP. We used the condition at 11 mW for 1 sec to irradiate R(n)s at stage 1, and R1+2.p at stage 3. About 30% of irradiated cells expressed GFP. Irradiated cells were sometimes damaged and failed to generate rays.

## RESULTS

### Background of the experiments: Development of rays and SMPs-PLX-1 signaling

Rays are sensory organs in the adult male tail, and play an important role during copulation (Figure 1(a), 3(a)). For readers who are not familiar with this structure, the development of epidermal cells in the male tail will be briefly summarized. Rays are generated from a row of epidermal cells called seam cells on the lateral side of the larval body (Emmons, 2005). Three bilateral pairs of seam cells, V5, V6 and T, in the posterior part of a male body produce the nine pairs of ray precursor cells, R1, R2-R6, R7-R9, at the third larval (L3) stage, respectively. (Figure 2(a)). A R(n) divides into two daughter cells, R(n).a and R(n).p (Figure 2(b), (i)), and then R(n).a generates three cells, which form a ray precursor cluster (Figure 2(c), (d), (e)) that eventually produces a single ray (n) (n=1 - 9). Meanwhile, R(n).ps, after shifting their position dorso-laterally to R(n).as, become enlarged, and fuse to each other. The cell-fusion occurs first between R1.p and R2.p (Figure 2(f), (g)), and then proceed posteriorly to eventually form a syncytial tail seam (set) (Figure 2(h)) (Sulston et al., 1980; Emmons, 2005). In this paper, for the convenience of description, we designate R(n)s and their descendants collectively as ray-lineage cells, and define the developmental stages as follows: stage 1: the stage prior to the cell division of R(n) (Figure 2(a)); stage 2: the stage after the cell division of R(n) and prior to fusion of R1.p and R2.p (Figure 2(b)-(f)); stage 3: the stage after the cell-fusion of R1.p and R2.p (Figure 2(g)). The cell generated by the fusion of R1.p and R2.p is referred to as R1+2.p in this paper (Figure 2(g)).

**Figure 1.**
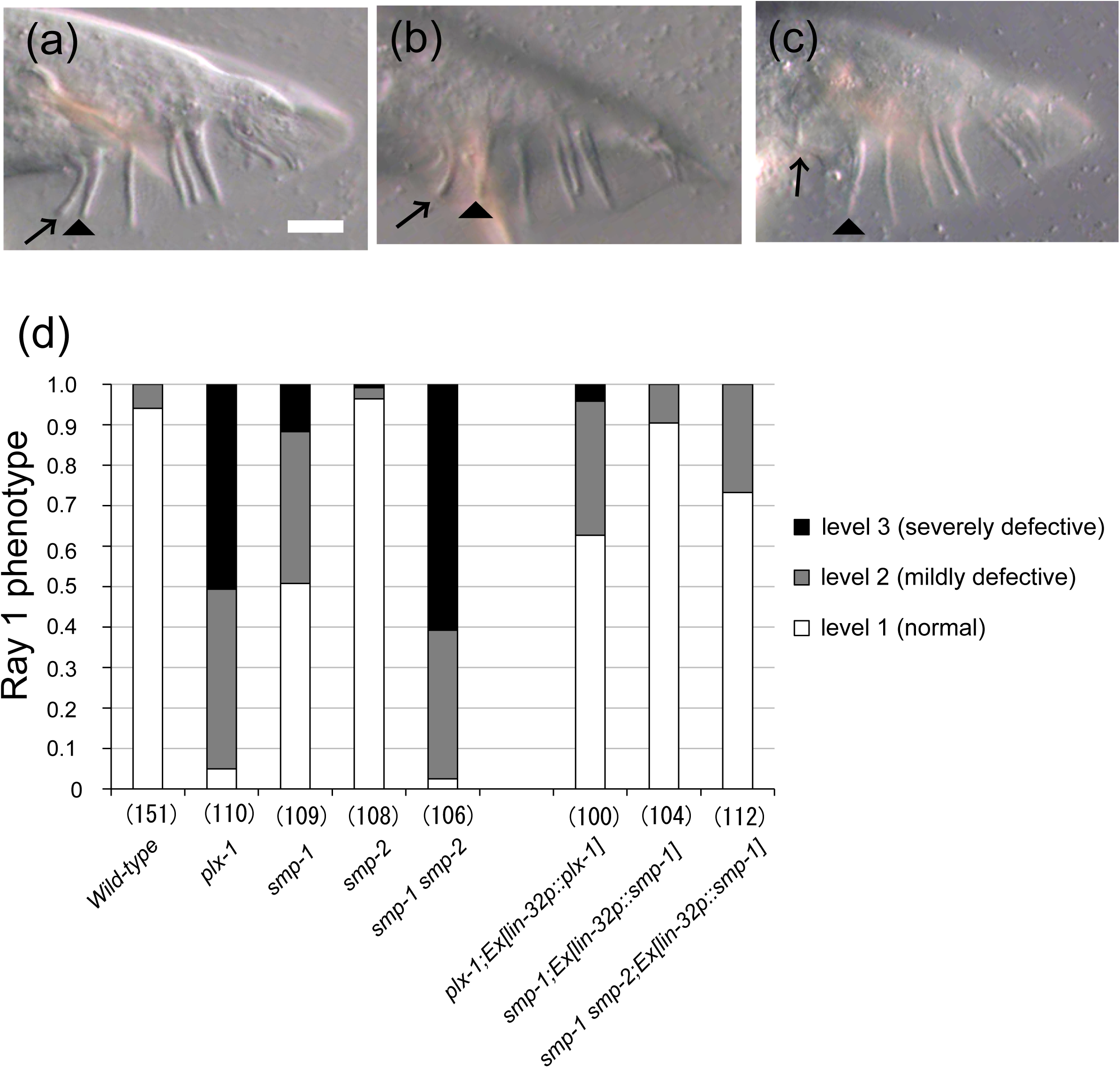
The male tail defect of *smp-1(ev715) smp-2(ev709)* and *plx-1(nc37)* mutants. (a) A lateral view of the tail of a wild-type adult male. Ray 1 (arrow) is normally located adjacent to the neighboring ray 2 (arrowhead)(Level 1). (b) Ray 1 is located inside the fan but is separated from ray 2 (Level 2). (c) The worm showed the severe displacement of ray 1, which is located outside of the fan (Level 3). (d) The penetrance of the ray 1 phenotype in each mutant. The *lin-32* promoter drives gene expression in R(n) cells and their descendants (ray-lineage cells). All worms carry the *him-5* mutation except for *smp-1; ncEx2038[lin-32p::smp-1]* and *smp-1 smp-2; ncEx2038[lin-32p::smp-1],* which carry the *him-8* mutation, to increase the incidence of male. Anterior is left, and dorsal is top. Scale Bar = 10 μm.

**Figure 2.**
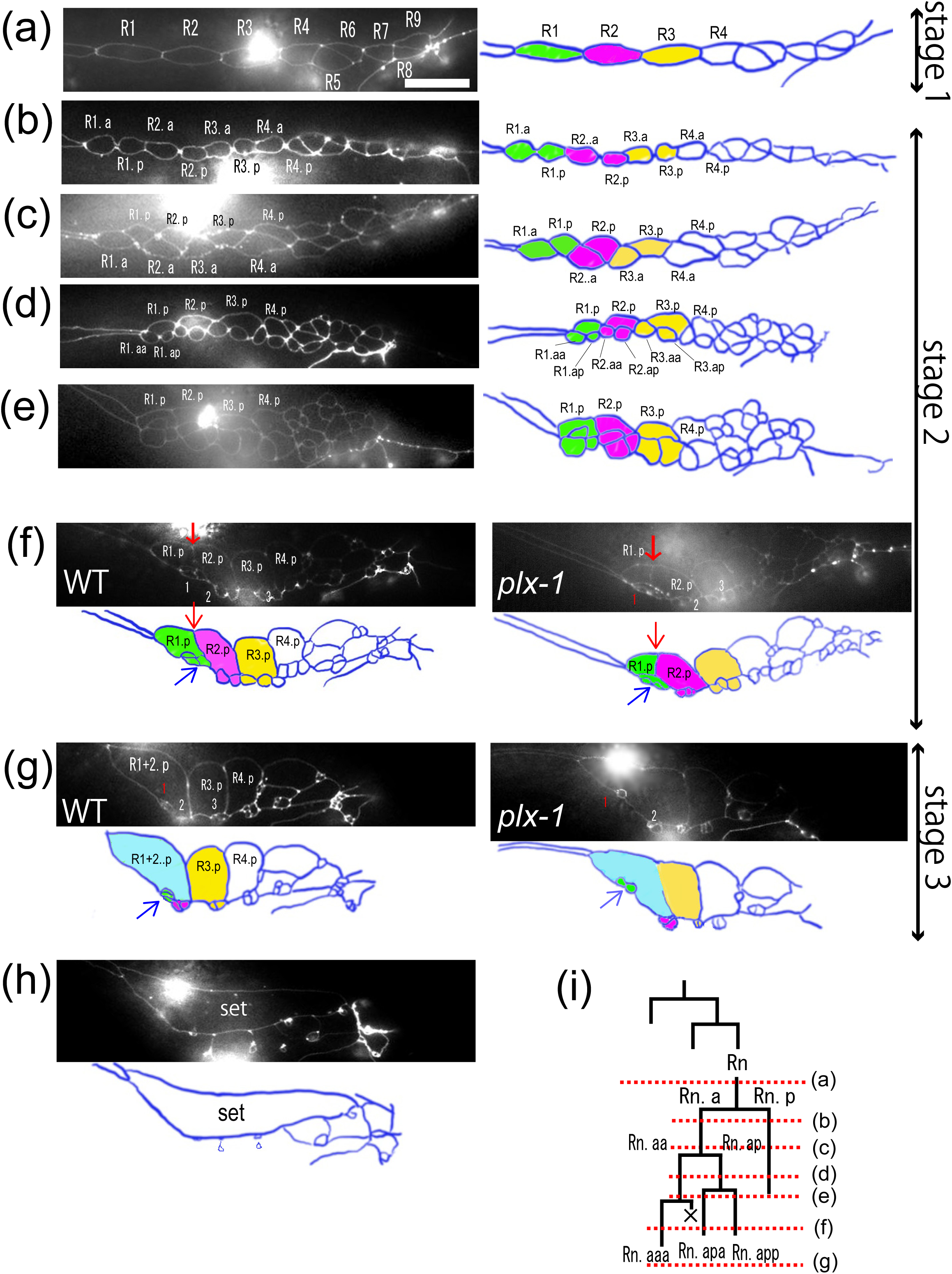
Development of lateral epidermis of the larval male tail. The apical border of epidermal cells was visualized with an apical junction marker, *AJM-1::GFP*. R(n) cells (a) and their descendants (b-e, h) in the wild-type male. The schematic drawing of each picture is shown on the right. R(1), R(2), and R(3) and their descendants are marked in green, magenta, and yellow, respectively. (f, g) The descendants of R(n) cells of a wild-type (WT) (left) and a *plx-1* mutant (right) male between the stages shown (e) and (h). The schematic drawing of each picture is shown below. (a) Nine R(n) cells are located on the lateral side of the larval male tail. (b) Each R(n) cell divides into two daughter cells, R(n).a and R(n).p. (c) The arrangement of R(n).a and R(n).p changes so that R(n).p is positioned dorsolateral to R(n).a. (d) R(n).a divides into two daughter cells, R(n).aa and R(n).ap. (e) R(n).aa and R(n).ap each divides again, producing R(n).aaa, R(n).aap, and R(n).apa, R(n).app, respectively. (f left) R(n).aap executes programmed cell death, while the remaining three cells, R(n).aaa, R(n).apa and R(n).app, form a ray precursor cluster. (Blue arrow indicates the ray precursor cluster 1). Then the apical surface of a ray precursor cluster shrinks while that of R(n).p enlarges. Red arrow indicates the apical junction between R1.p and R2.p. (g) R1.p fuses with R2.p to form R1+2.p (light blue). (h) After formation of R1+2.p, the remaining R(n).ps gradually fuse together to form a syncytial tail seam (set) (f,g right) In a *plx-1* mutant male, the border between R1.p and R2.p shifts anteriorly (red arrow in f), and the position of the ray precursor cluster 1 (blue arrow in g) is dislocated anteriorly. (i) Cell lineages of R(n) and its descendant cells. ×: programmed cell death. Anterior is left, and dorsal is top. Scale Bars = 10 μm.

**Figure 3.**
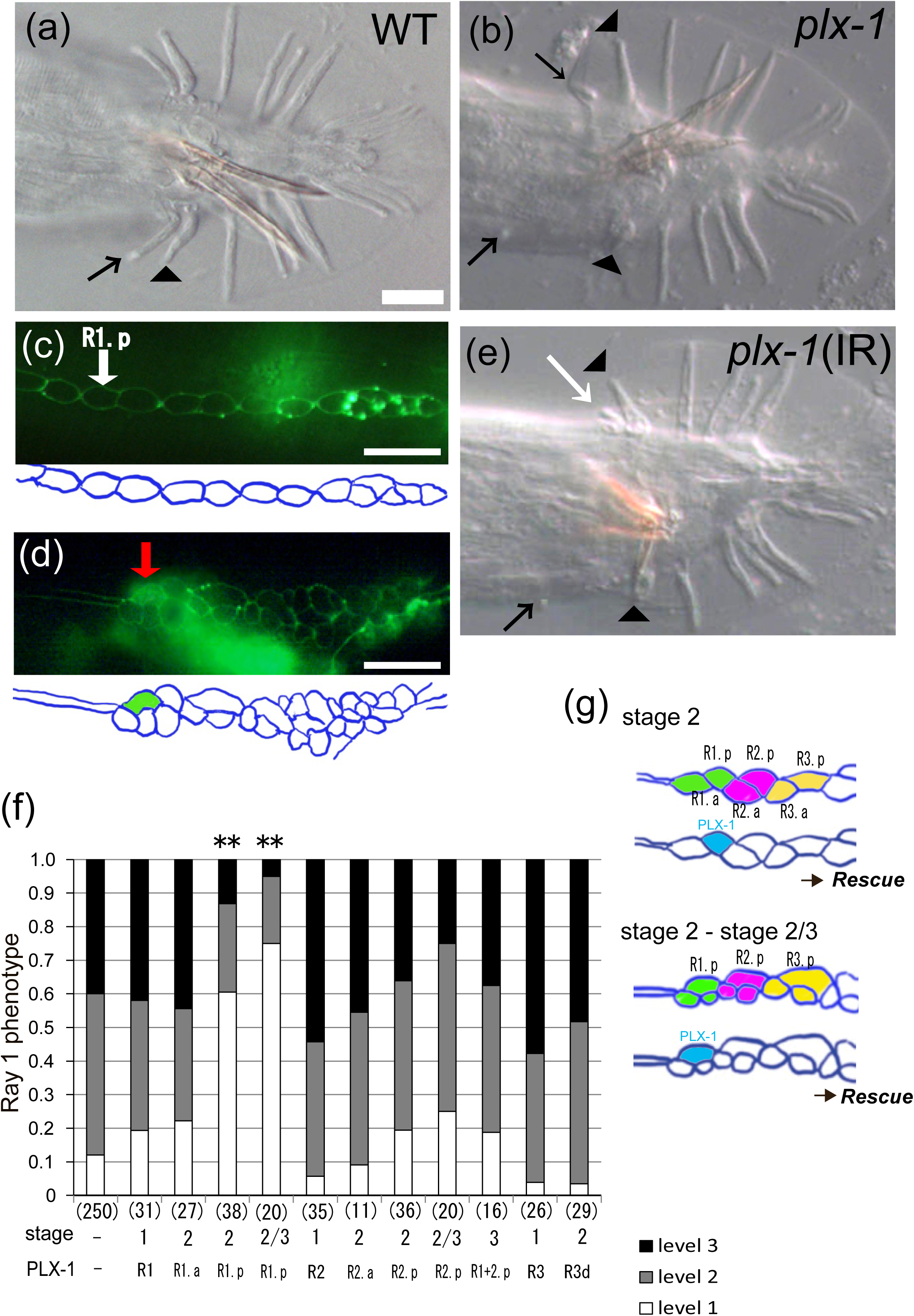
Induced expression of PLX-1 in R1.p efficiently rescued the ray 1 defect in *plx-1* mutants. A DIC photomicrograph of the ventral view of adult rays of (a) a wild-type and (b) a *plx-1* mutant animal. (a) Ray 1 (arrow) is juxtaposed to ray 2 (arrowhead). (b) In the *plx-1* mutant, ray 1 (arrows) on the right side (Level 3) and on the right side (Level 2) were both shifted anteriorly, separated from ray 2 (arrowheads). (c) R1.p cell (white arrow) of a *plx-1; ncIs19[hsp::plx-1; hsp::gfp; rol-6(su1006)]; ncIs13[ajm-1::gfp]* worm was IR-irradiated. (d) Three hours after irradiation, the expression of GFP, an induction marker, was found only in the cytoplasm of the targeted R1.p (red arrow). A diffuse green signal in the area below ray-lineage cells is due to autofluorescence. (e) Induced expression of PLX-1 in the left R1.p at stage 2 rescued the ray 1 defect on the irradiated side (white arrow). Arrows indicate ray 1, and arrowheads ray 2 (a, b, e). Scale Bars = 10 μm. (f) The ray 1 phenotype in the adult tail of IR-irradiated worms. *plx-1; ncIs19[hsp::plx-1; hsp::gfp; rol-6(su1006)]; ncIs13[ajm-1::gfp]* worms were used. The number of worms that are successfully induced and scored are shown in the parenthesis. “stage - / PLX-1 –“ indicates the ray 1 phenotype of non-irradiated worms. Stage“2/3” indicates the cases where worms that were irradiated at stage 2 had developed into stage 3 within three hours after irradiation. “R3d” indicates the combined scores of induction in R3 daughter cells, R3.a and R3.p. Induced expression of PLX-1 in R1.p at stages 2 and 2/3 rescued the ray 1 defect (** p<0.01, Ryan’s method for multiple comparison for the proportion of the Level 1 phenotype). (g) Summary of induced expression of PLX-1. Induction in R1.p (blue) rescued the ray 1 defect.

In the majority of wild-type animals, the anterior most ray 1 is juxtaposed to the neighboring ray 2 (Figure 1(a), 3(a)), which we define as “Level 1” for the position of ray 1. Since the position of the ray precursor cluster 1 in *plx-1* and *smp-1 smp-2* mutants shifts anteriorly compared to that in wild-type animals at the L4 stage (Figure 2(g)) (Fujii et al., 2002; Nukazuka et al., 2008), the mutant adult males show the highly penetrant defect in which the position of ray 1 is displaced anteriorly. In many cases, ray 1 is either located outside of the fan (Level 3) (Figure 1(c), the right side of the tail in Figure 3(b)) or inside of the fan but is separated from ray 2 (Level 2) (Figure 1(b), the left side of the tail in Figure 3(b)). PLX-1 is expressed in the all ray-lineage cells (Fujii et al., 2002) and the defective positioning of ray 1 in *plx-1* mutants was rescued by expression of a PLX-1 cDNA under the control of the *lin-32* promoter (Figure 1(d)), which drives expression of transgenes in the all ray-lineage cells (Portman and Emmons, 2000), as reported previously (Nakao et al., 2007; Nukazuka et al., 2008). Similarly, the defective positioning of ray 1 in *smp-1 smp-2* mutants was rescued by expression of a SMP-1 cDNA under the control of the *lin-32* promoter with the transgene *ncEx2038[lin-32p::smp-1; rol-6(su1006)]* (Figure 1(d)).

While these rescue experiments indicate that SMPs-PLX-1 signaling among ray-lineage cells determines the proper position of ray 1, the cells that send and/or receive the signal remain to be specified. To examine further where and when the SMPs signal is required, we induced to express SMP-1 and PLX-1 in a single cell of each mutant animal by using IR-LEGO. For induction of PLX-1 and SMP-1, we used the transgenes, *ncIs19[hsp::plx-1; hsp::gfp]* and *ncIs24[hsp::smp-1; hsp::gfp]*, which contains *hsp::plx-1* and *hsp::smp-1*, respectively, and *hsp::gfp* as an induction marker. In this experiment, we irradiated a ray-lineage cell at different developmental stages (Figure 3(c)) and checked the GFP expression 3 hours after irradiation (Figure 3(d)). Then, one day after irradiation we scored the position of ray 1 (Figure 3(e)).

### Induction of PLX-1 in R1.p rescued the ray 1 defect of *plx-1* mutants

First we tried to express PLX-1 in R1 (a V5 descendant) of the *plx-1; ncIs19[hsp::plx-1; hsp::gfp]* worm at stage 1. In worms that successfully expressed PLX-1 in R1, the proportion of each level of the ray 1 position did not differ significantly from that in *plx-1* worms (n=31. Level 1: 20%, Level 3: 42%) (Figure 3(f)), indicating that the ray 1 displacement defect was not rescued. Expression of PLX-1 in the other R(n)s at stage 1 also did not rescue the ray 1 defect of *plx-1* mutants (Level 1: 6% n=35, 4% n=26, 7% n=14 for induction in R2, R3 and R4 respectively) (Figure 3(f)).

Then worms were irradiated at stage 2, when R(n) had already completed cell division to give rise to R(n).a and R(n).p. Induction of PLX-1 expression in R1.p of *plx-1* mutants rescued the ray 1 defects significantly: the proportion of Level 1, which represents normal positioning (Figure 3(e)), increased to more than 60% (n=38, p<0.01) while that of Level 3, which represents severe anterior displacement, decreased to less than 10% (Figure 3(f)). In contrast, induction of PLX-1 expression in the other cells at stage 2 did not lead to significant rescue of the ray1 defects. Induction of GFP alone, which serves as a control of IR-irradiation, in R1.p of *plx-1* mutants did not affect the position of ray 1 (Figure S1).

When we irradiated R1.p or R2.p at stage 2, we occasionally found the GFP expression in R1+2.p as a result of fusion of R1.p with R2.p by 3 hours after irradiation. Such cases will be referred to induction at “stage 2/3” in this paper. Induction of PLX-1 expression in R1.p at stage 2/3 also rescued the ray 1 defect significantly (Level 1: 75% n=20, p<0.01) (Figure 3(f)). On the other hands, induction in R2.p at stage 2/3 did not rescue the ray 1 defect significantly (n=20). When worms were irradiated at stage 3, induction of PLX-1 expression in R1+2.p or other cells (data not show) did not rescue the ray 1 defect.

These results indicate that function of PLX-1 in R1.p at stage 2 or 2/3 is required for proper positioning of ray 1 (Figure 3(g)).

### Induction of SMP-1 in R1.p, R2.a or R2.p rescued the ray 1 defect of *smp-1 smp-2* mutants

Next, we tried to examine the functional site of the semaphorin ligand involved in positioning of ray 1. While two membrane-bound semaphorins, SMP-1 and SMP-2, both bind to PLX-1, and are involved in regulation of the positioning of ray 1 (Ginzburg et al., 2002; Fujii et al., 2002), SMP-1 has a dominant role; In *smp-2* single mutants the ray 1 displacement defect is almost negligible, and expression of SMP-1 cDNA under the control of the *lin-32* ray-lineage specific promoter is sufficient for rescuing the ray 1 defect of *smp-1 smp-2* double mutants nearly completely (Figure 1(d)). Therefore, we focused on SMP-1 and induced it in a single ray-lineage cell in *smp-1 smp-2*; *ncIs24[hsp::smp-1; hsp::gfp]* worms by using IR-LEGO (Figure 4(a)). The induced expression of SMP-1 in ray-lineage cells at stage 1 or stage 3 did not rescue the phenotype of ray 1 displacement. Expression of SMP-1 in a single R1.p, R2.a or R2.p at stage 2 rescued the ray 1 defect (Level 1: 50% n=34, 53% n=15, and 53% n=19, for induction in R1.p, R2.a or R2.p, respectively. p<0.05). Induction of GFP alone in these cell of *smp-1 smp-2* mutants did not affect the position of ray 1 significantly (Figure S2), confirming that expression of SMP-1 in either R1.p, R2.a or R2.p is required for proper positioning of ray 1 (Figure 4(b)). In another control experiment, induced expression of PLX-1 in R1.p of *smp-1 smp-2* mutants did not affect the ray 1 phenotype (Level 1: 4% n=69).

**Figure 4.**
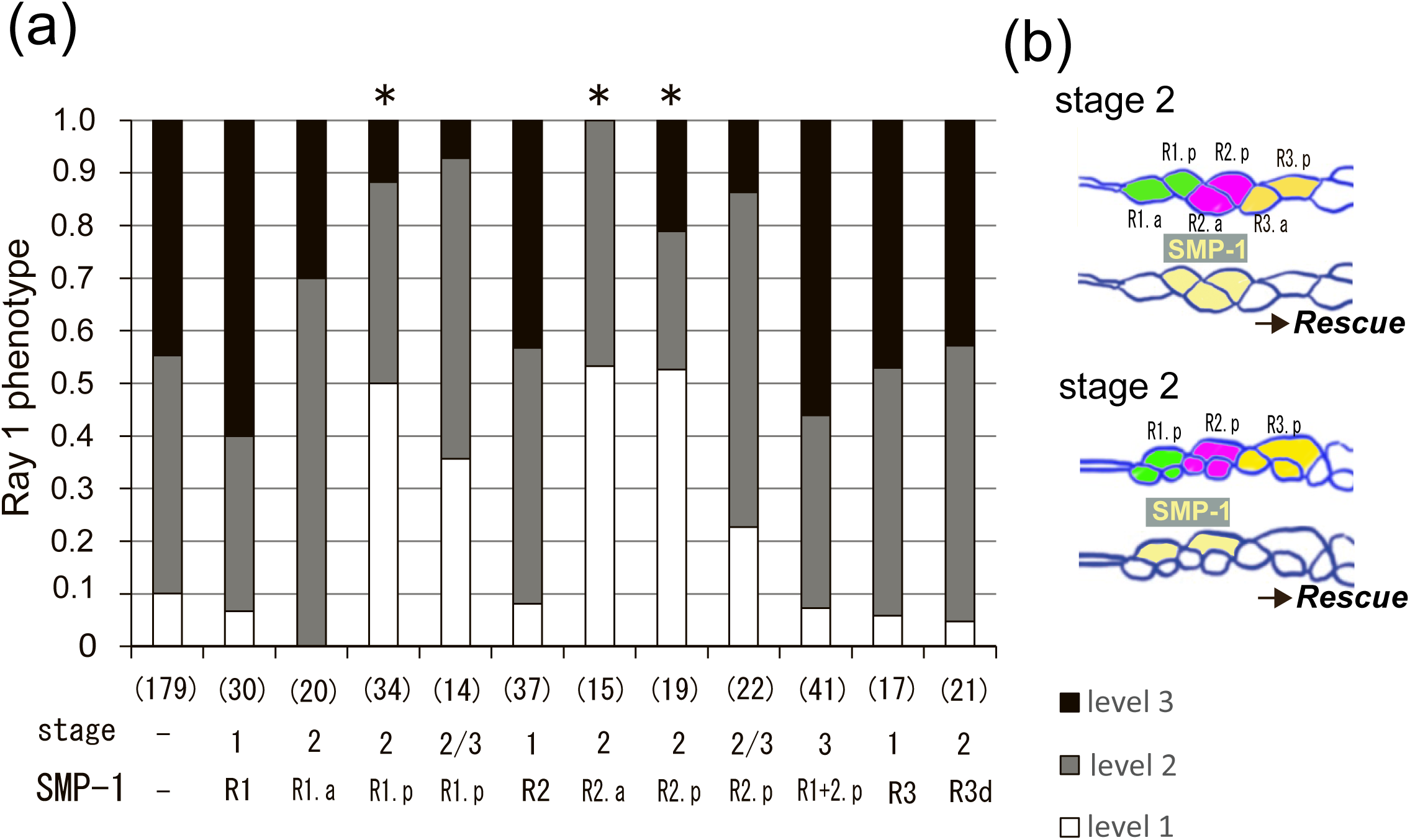
Induced expression of SMP-1 in either R1.p, R2.a or R2.p efficiently rescued the ray 1 defect in *smp-1 smp-2* mutants. (a) The ray 1 phenotype in the adult tail of IR-irradiated *smp-1 smp-2; ncIs24[hsp::smp-1; hsp::gfp; rol-6(su1006)]; ncIs13[ajm-1::gfp]* worms was scored. SMP-1 was induced in each R(n) cell and its daughters of *smp-1 smp-2* double mutants. “R3d” indicates the combined scores of induction to R3 daughter cells, R3.a and R3.p. The number of worms that are successfully induced and scored are shown in the parenthesis. Induced expression of SMP-1 in R1.p, R2.a or R2.p at stage 2 rescued the ray 1 defect (* p<0.05, Ryan’s method for multiple comparison for the proportion of the Level 1 phenotype). (b) Summary of induced expression of SMP-1. Induction in one of the pale yellow cells rescued the ray 1 defect.

On the other hand, expression of SMP-1 in R1.a or daughter cells of R3 at stage 2 did not rescue the ray 1 displacement defect (Level 1: 0% n=20 and 5% n=21 for induction in R1.a and R3d, respectively). In the case of induction at stage 2/3, the proportion of Level 1 and Level 3 appeared to be increased and reduced, respectively, by expressing SMP-1 in either R1.p or R2.p, (Level 1: 36% n=14 for R1.p; 23% n=22 for R2.p), though the results are not considered to be statistically significant.

As shown in the preceding section, expression of PLX-1 in R1.p appears critical for proper positioning of ray 1. Both R2.a and R2.p, in which induction of SMP-1 rescued the ray 1 defect, are positioned posterior to and in contact with R1.p during a certain period of the development of the male tail (Figure 2, 6(c), (d)). In contrast to this, induced expression of SMP-1 in R1.a, which is in contact with but positioned anterior to R1.p, failed to restore proper positioning of ray 1. These results suggest that activation of SMPs-PLX-1 signaling in the posterior part of R1.p via the adjacent cells is critical for proper positioning of ray 1, presumably by shifting the position of the border between R1.p and R2.p posteriorly. Therefor we propose a model, in which SMP-1 on R2.a or R2.p promotes or permits the extension of the contacting edge of the neighboring R1.p (Figure 6(c), (d), (f)).

### Co-induction of SMP-1 and PLX-1 in R1.p rescued the ray 1 defect of *smp-1 smp-2*; *plx-1* mutants partly

The model described above, however, does not explain another finding that induced SMP-1 in R1.p also rescued the ray 1 defect. We think that this apparent contradiction could be resolved by assuming the *cis*-interaction between SMP-1 and PLX-1. Our previous study showed that SMP-1, which was overexpressed in a VPC by using IR-LEGO, interacts with co-expressed PLX-1 in *cis* and triggers the signal downstream to PLX-1 (Suzuki et al., 2022). Similarly, the present finding that SMP-1 expression in R1.p rescues the ray 1 defect in *smp-1* mutants might be explained if induced SMP-1 *cis*-activates PLX-1 on the same R1.p cell. To examine this possibility, we induced the expression of both SMP-1 and PLX-1 in a single ray-lineage cell in *smp-1 smp-2; plx-1* triple mutants background by using *smp-1 smp-2; plx-1*; *ncIs210[hsp::plx-1; hsp::smp-1; hsp::gfp; rol-6(su1006)]* worms. We focused our analysis on the effect of induction in R1.a, R1.p, R2.a or R2.p at stage 2. We found that IR-irradiation to R1.p rescued the ray 1 displacement defect in 22% of the *smp-1 smp-2; plx-1* mutant worms (n=22 p<0.05.). IR-irradiation to the other cells did not affect the ray 1 phenotype (Level 1: less than 5% n=54) (Figure 5(a)). These data are consistent with the idea that SMP-1 induced by IR-LEGO *cis*-activates PLX-1 in R1.p (Figure 6(e), (g)).

**Figure 5.**
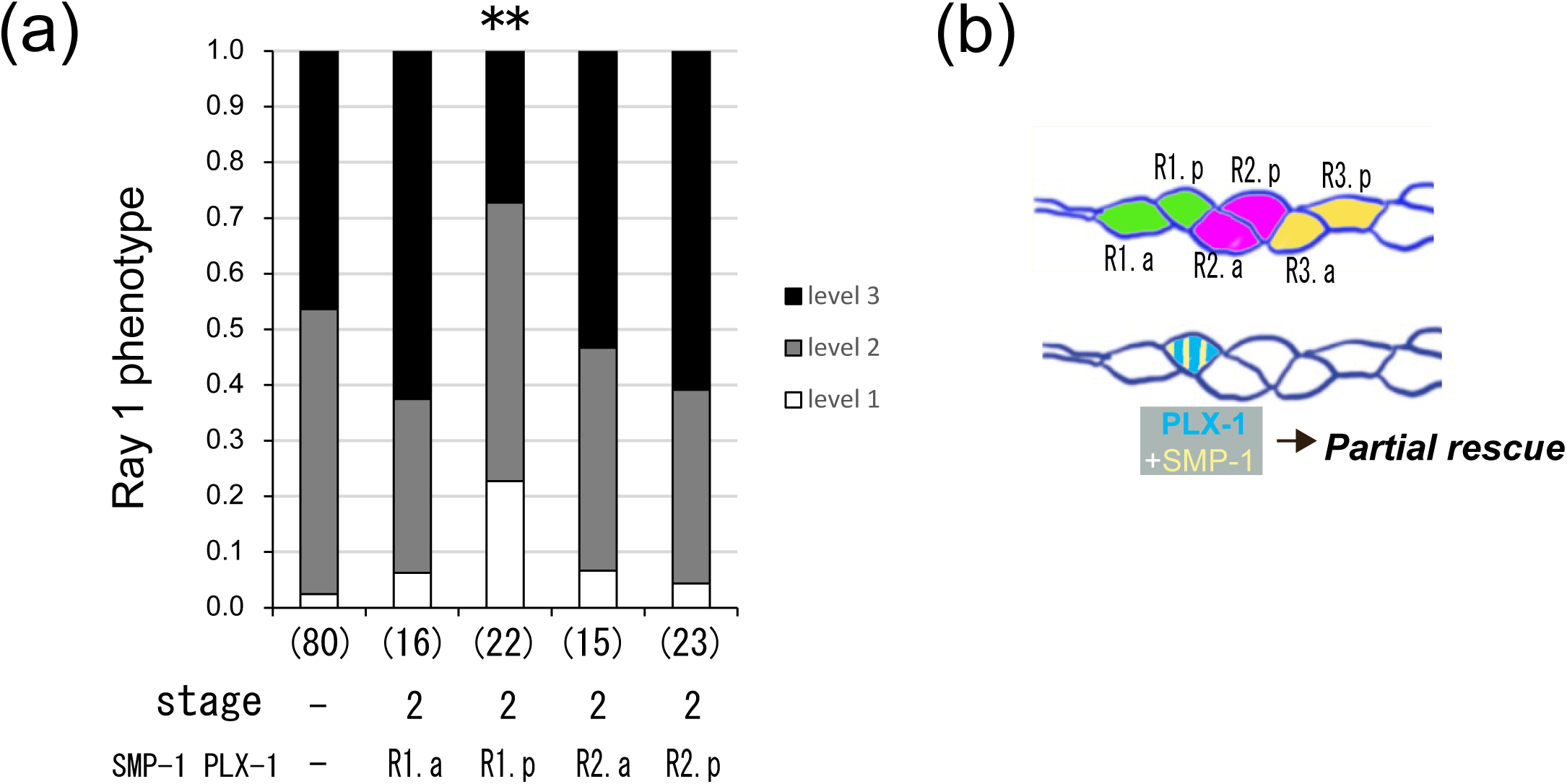
Induce expression of SMP-1 and PLX-1 in *smp-1 smp-2; plx-1* mutants of single R1.p partly rescued the ray 1 displacement. (a) *smp-1 smp-2; plx-1; ncIs210 [hsp::plx-1; hsp::smp-1; hsp::gfp; rol-6(su1006)]; ncIs13[ajm-1::gfp]* worms at stage 2 were used in this experiment (p<0.01, Ryan’s method for multiple comparison for the Level 1 phenotype). The total number of IR-irradiated worms are shown in the parenthesis. (b) Summary of induced expression of SMP-1 and PLX-1 in *smp-1 smp-2; plx-1* mutants.

**Figure 6.**
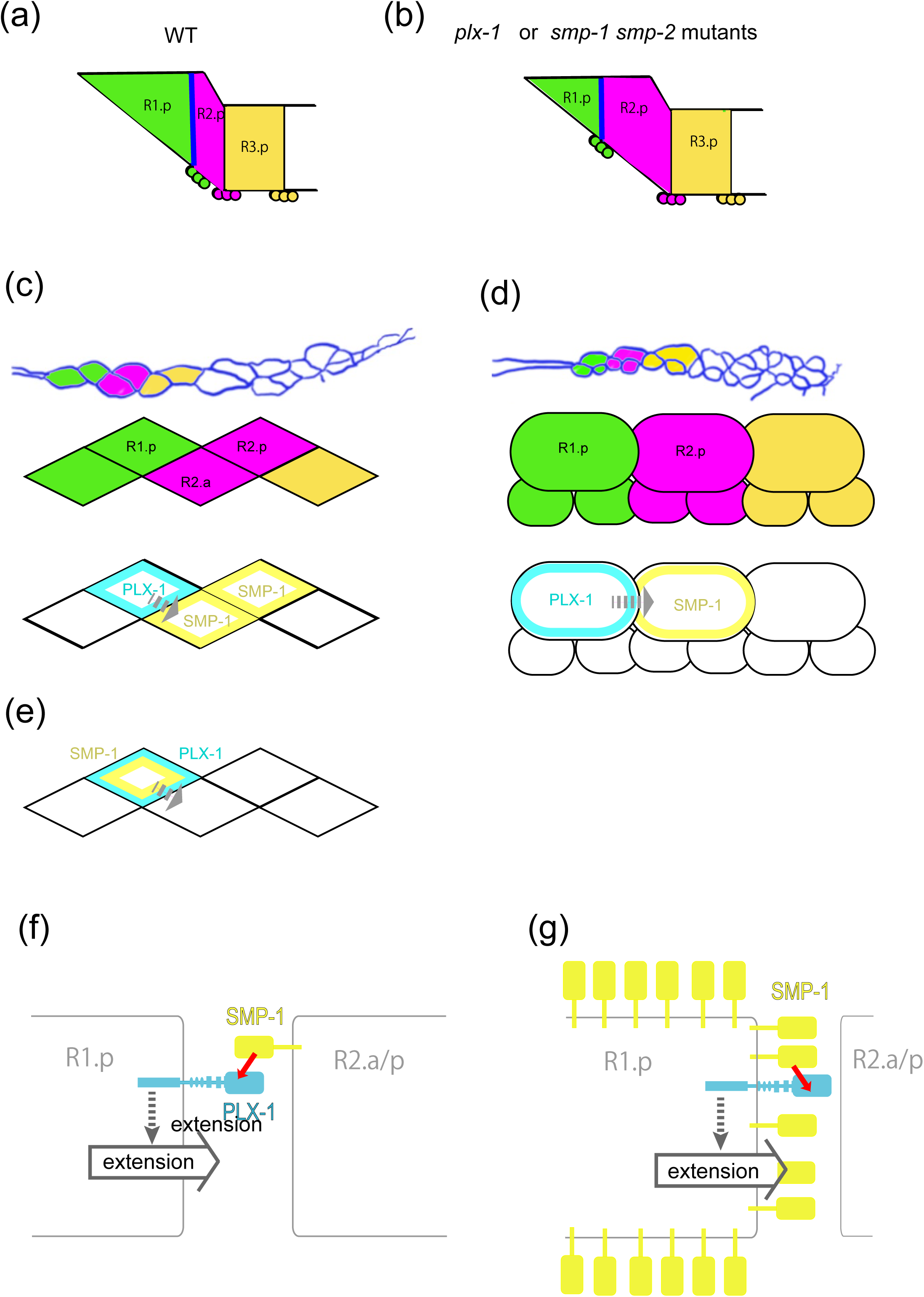
A model of SMPs signaling among ray-lineage cells. (a, b) Schema of arrangement of R1.p, R2p and the ray precursor cluster 1 in a wild-type (a) and a *plx-1* or *smp-1 smp-2* mutant (b) animal. The position of the ray precursor cluster 1 shifts anteriorly in the mutant. The blue line shows the apical border between R1.p and R2.p, which determine the position of the ray precursor cluster 1. (c-e) Summary of the rescue experiments by using IR-LEGO. Expression of PLX-1 in R1.p and SMP-1 in R2.a/p (c, d) is sufficient for rescuing the ray 1 defect in mutants, suggesting that the *trans*-interaction between SMP-1 and PLX-1 (f) promotes extension of the posterior part of R1.p. Induced expression of PLX-1 and SMP-1 in R1.p (e) also partly rescued the ray 1 defect in the mutants, suggesting that the *cis*-interaction between SMP-1 and PLX-1 (g) can promotes the extension of R1.p.

## DISCUSSION

By using IR-LEGO, we have successfully induced the expression of functional genes in a single epidermal cell of the larval male tail of *C. elegans*. We showed that expression of PLX-1 in R1.p is sufficient for rescuing the ray 1 displacement defect in *plx-1* mutants. We also showed that the SMPs signal emanating from R2.p or R2.a, the cells located adjacent to and posterior to R1.p during a certain period of development, is sufficient for positioning ray 1 properly. Although a previous study suggested that the SMPs signal emanating from remote cells affects ray-lineage cells (Dalpé et al., 2004), the present results suggest that SMPs signal mediated in a cell-cell contact-dependent manner is sufficient for determining the position of ray 1.

It was suggested that the position of a ray precursor cluster, which later generate a ray, is determined by the position of the border between two adjacent R(n).p cells and Hyp 7; the ray precursor cluster appears retained at the position where the surface of the three cells intersect (Baird et al., 1991) (Figure 6(a)). Accordingly, our previous studies confirmed that the ray precursor cluster 1 is localized at the junction between R1.p, R2.p, and Hyp 7 in both WT and *plx-1* worms (Fujii et al., 2002; Nukazuka et al., 2008). Thus, the anterior displacement of ray 1 in *plx-1* mutants is attributable to the shift of the border between R1.p and R2.p anteriorly to its normal position (Figure 6(b)). Our finding is consistent with a model in which the SMPs signal emanating from R2.p and R2.a acts on, and promotes or permits the extension of the posterior edge of R1.p to keep the border between R1.p and R2.p at a proper posterior position (Figure 6(f)).

Induction of PLX-1 expression at stage 2 or 2/3, when the shape of R1.p and R2.p changes dramatically, rescued the ray 1 defects strongly. Earlier induction of PLX-1 or SMP-1 at stage 1 failed to rescue the defects in the *plx-1* and *smp-1 smp-2* mutants, respectively. It might be that the proteins induced at stage 1 are no longer present by stage 2.

Unexpectedly, we found that induced expression of SMP-1 in R1.p, the cell on which we suggest the SMPs signal acts, efficiently rescues the ray 1 positioning defect in *smp-1 smp-2* mutants. We also found that induction of both SMP-1 and PLX-1 in R1.p of *smp-1 smp-2; plx-1* mutants partly rescues the ray 1 defect. These findings suggest that SMP-1, a cell membrane-bound protein, induced by IR-LEGO *cis*-activates PLX-1 which is co-expressed on R1.p (Figure 6(g)). The *cis*-interaction between SMP-1 and PLX-1 was reported previously (Mizumoto and Shen, 2012). Our previous study also showed that SMP-1 *cis*-activates PLX-1 in VPCs only when SMP-1 was overexpressed; when being expressed at a physiological level, SMP-1 failed to *cis*-activate PLX-1 (Suzuki et al., 2022). Although it is likely that induction with IR-LEGO leads to overexpression of SMP-1 in R1.p, whether SMP-1 expressed at a physiological level on R1.p can *cis*-activates PLX-1 remains to be determined.

The results suggest that SMPs signaling promotes or permits, rather than inhibits, the extension of a cell. This, however, is in contrast to our previous finding on VPCs that SMPs signaling act as a cell contact-dependent stop signal to arrest the extension of neighboring cells. It is not clear why the consequence of activation of SMPs signaling differs between these two types of *C. elegans* epidermal cells. It is also intriguing that SMPs-PLX-1 signaling mutants do not exhibit apparent morphological defects in ray-lineage cells other than R1.p, although our previous studies indicated that SMPs signaling is activated in other ray-lineage cells in addition to R1.p (Nukazuka et al., 2008; Tanaka et al., 2020). What cellular mechanisms confer R1.p the specific morphological response to SMPs signaling remains to be investigated. In the vertebrates, it is well known that some semaphorins act not only as repulsive but also as attractive axon guidance cues depending on the situation (Song et al., 1998; Falk et al., 2005). Therefore, context-dependence of output on cell shape appears to be a general feature of the semaphorin signaling systems.

## ACKNOWLEDGEMENTS

We thank professor Yoichi Oda and the members of the Oda Laboratory for discussion and comments, Hiromichi Omiya of Sigma Koki Co. Ltd for technical supports, Yuji Kohara for cDNA clones, Andrew Fire for pPD29.26 and pPD49.78, and Joe Culotti and Jeff Shimske for strains. Some strains were provided by the Caenorhabditis Genetic Center, which is funded by the National Institute for Health National Center for Research Resources (P40 OD010440). This work was supported by grants from the Ministry of Education, Science, Sports Culture and Technology (MEXT), Japan (S.T.), and a grant from The Naito Memorial Foundation (S.T.). M.S. is a Doctoral Course (DC1) Research Fellow of the Japan Society for the Promotion of Science.

## Supplementary materials

**Supplementary Figure S1.**
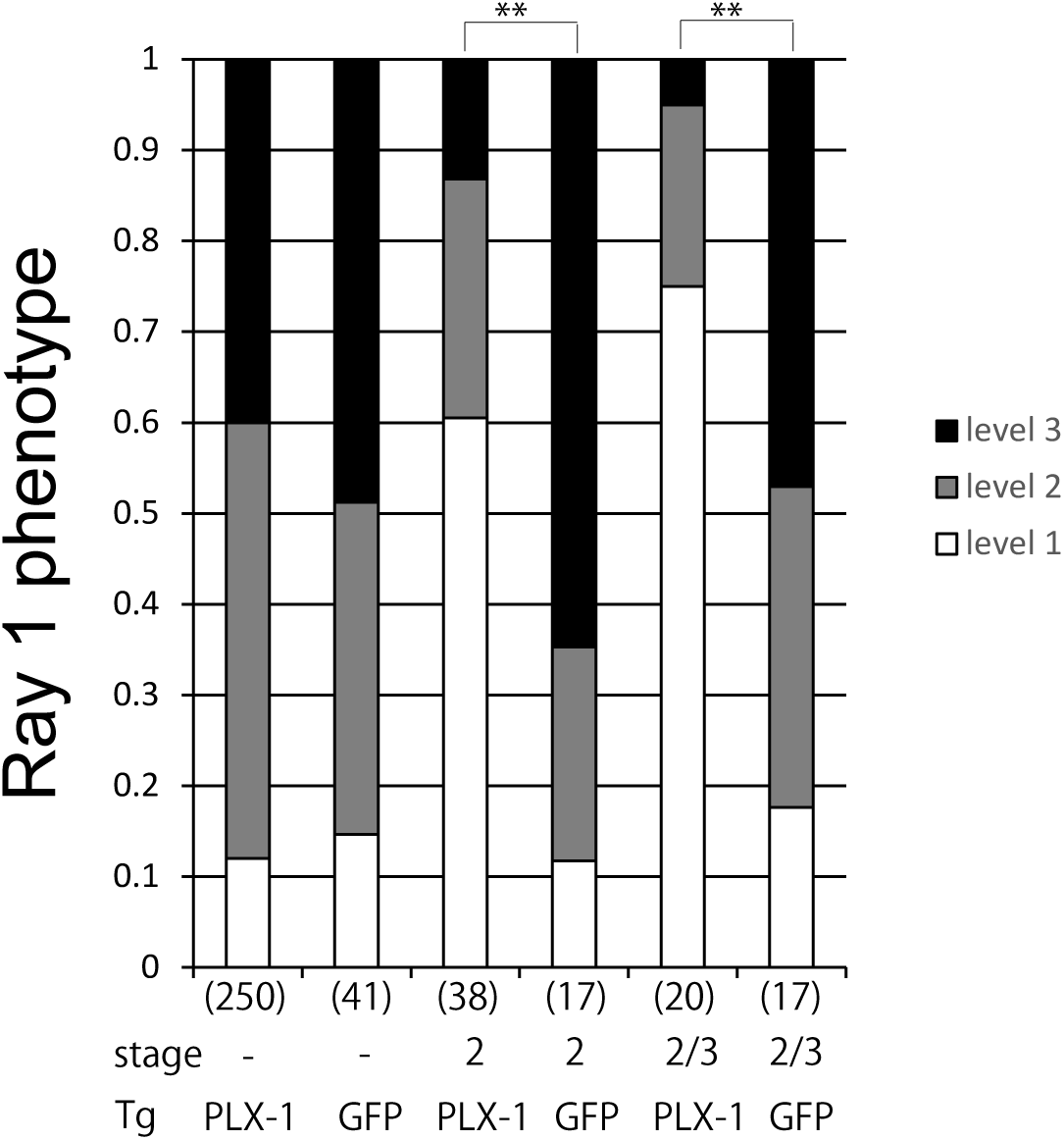
(G) The ray 1 phenotype in the adult tail of IR-irradiated *plx-1; ncIs19[hsp::plx-1; hsp::gfp]; ncIs13[ajm-1::gfp]* worms compared with that of *plx-1; jcIs1[ajm-1::gfp; rol-6(su1006)]*; *ncIs211[hsp::gfp]* worms that serve as controls. R1.p was targeted for gene induction at stage 2 and stage 2/3. Induced expression of GFP alone in control worms did not affect the ray phenotype (** p<0.01, Fisher’s exact test).

**Supplementary Figure S2.**
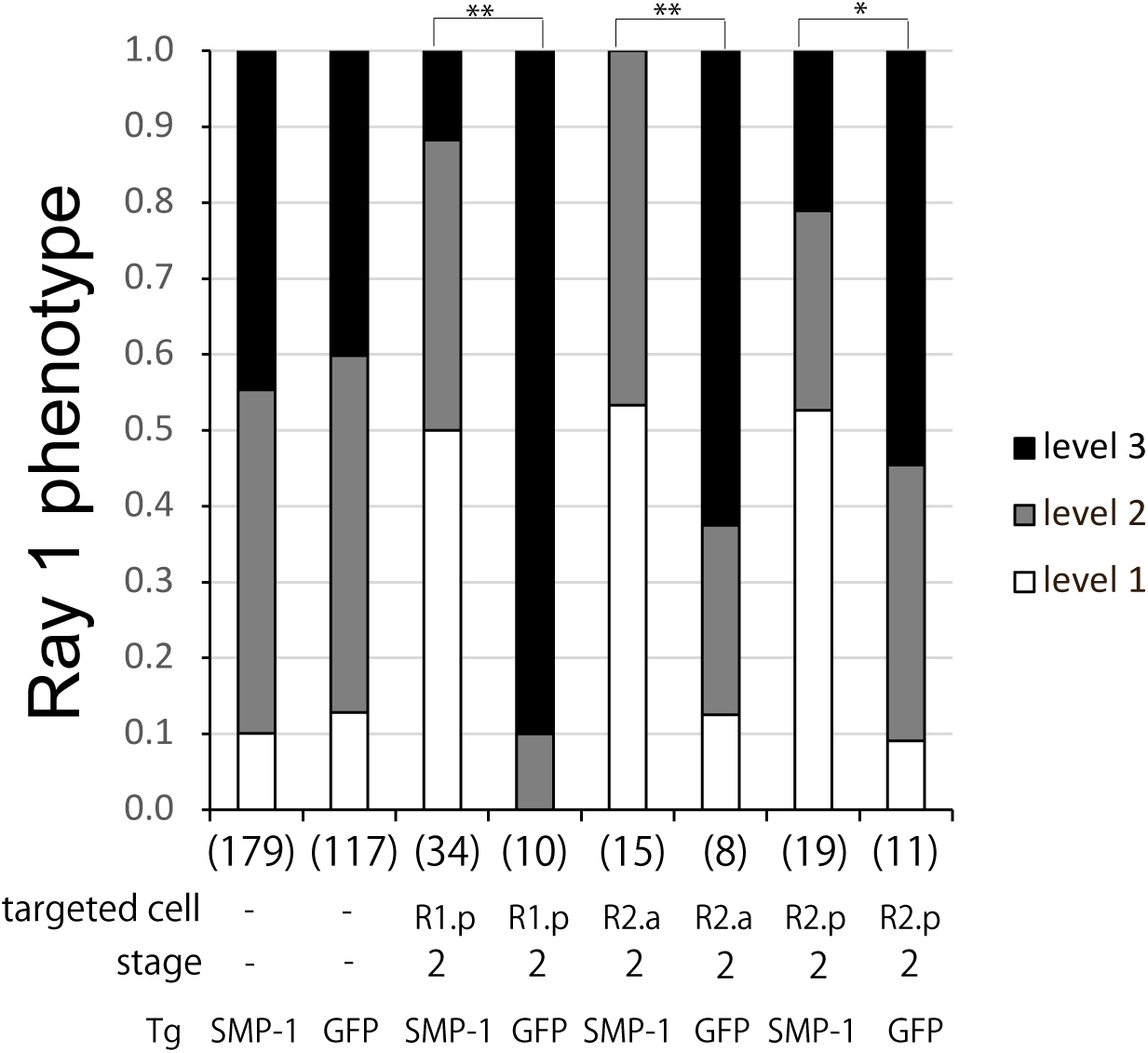
The ray 1 phenotype in the adult tail of IR-irradiated *smp-1 smp-2; ncIs24[hsp::smp-1; hsp::gfp]; ncIs13[ajm-1::gfp]* worms compared with that of *smp-1 smp-2; ncIs13[ajm-1::gfp]; ncIs17[hsp::gfp]* worms that serve as control. Induced expression of GFP alone in R1.p, R2.a or R2.p of control worms did not affect the ray phenotype. (** p<0.01, * p<0.05, Fisher’s exact test).

**Supplementary Table 1.**
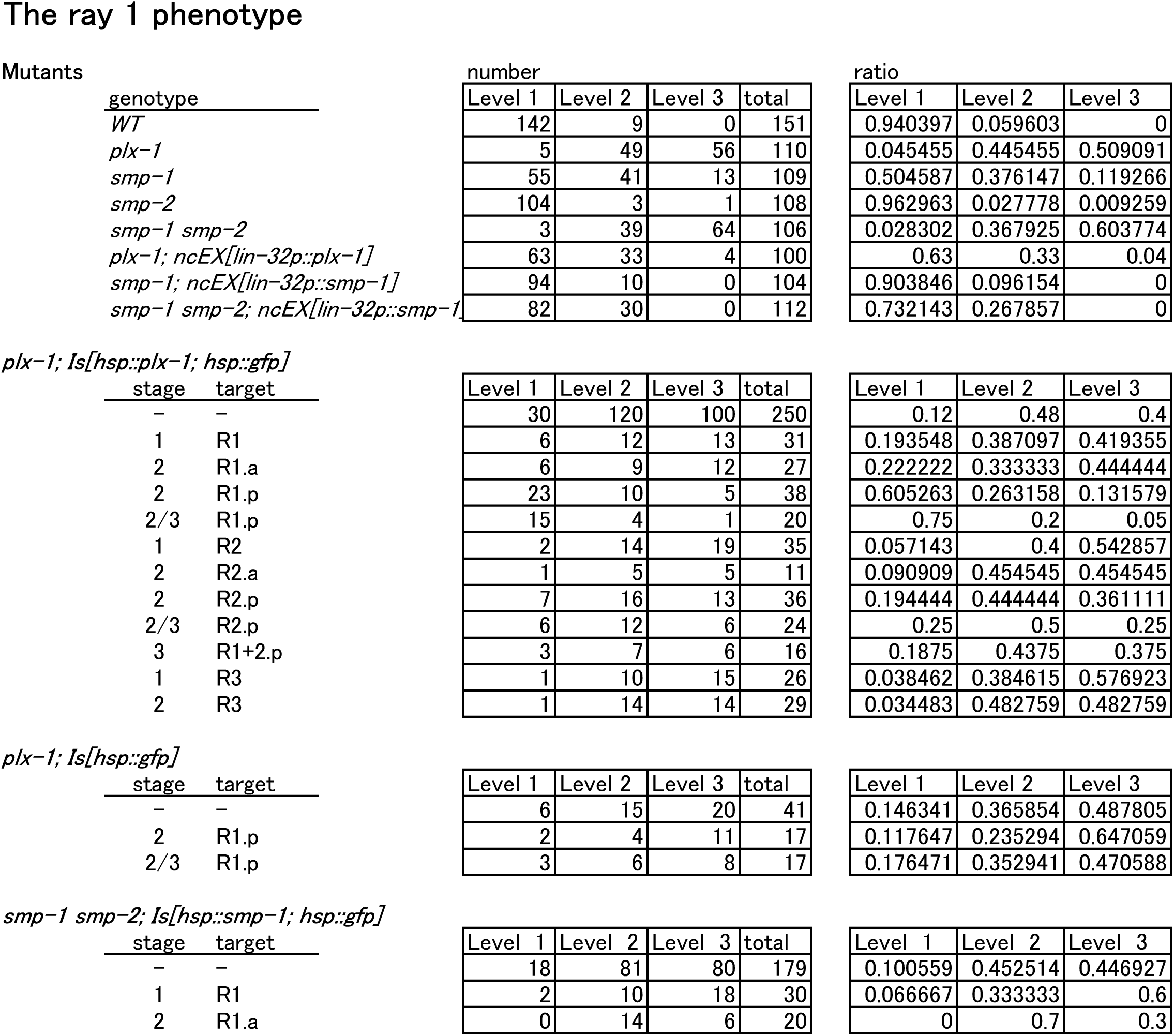
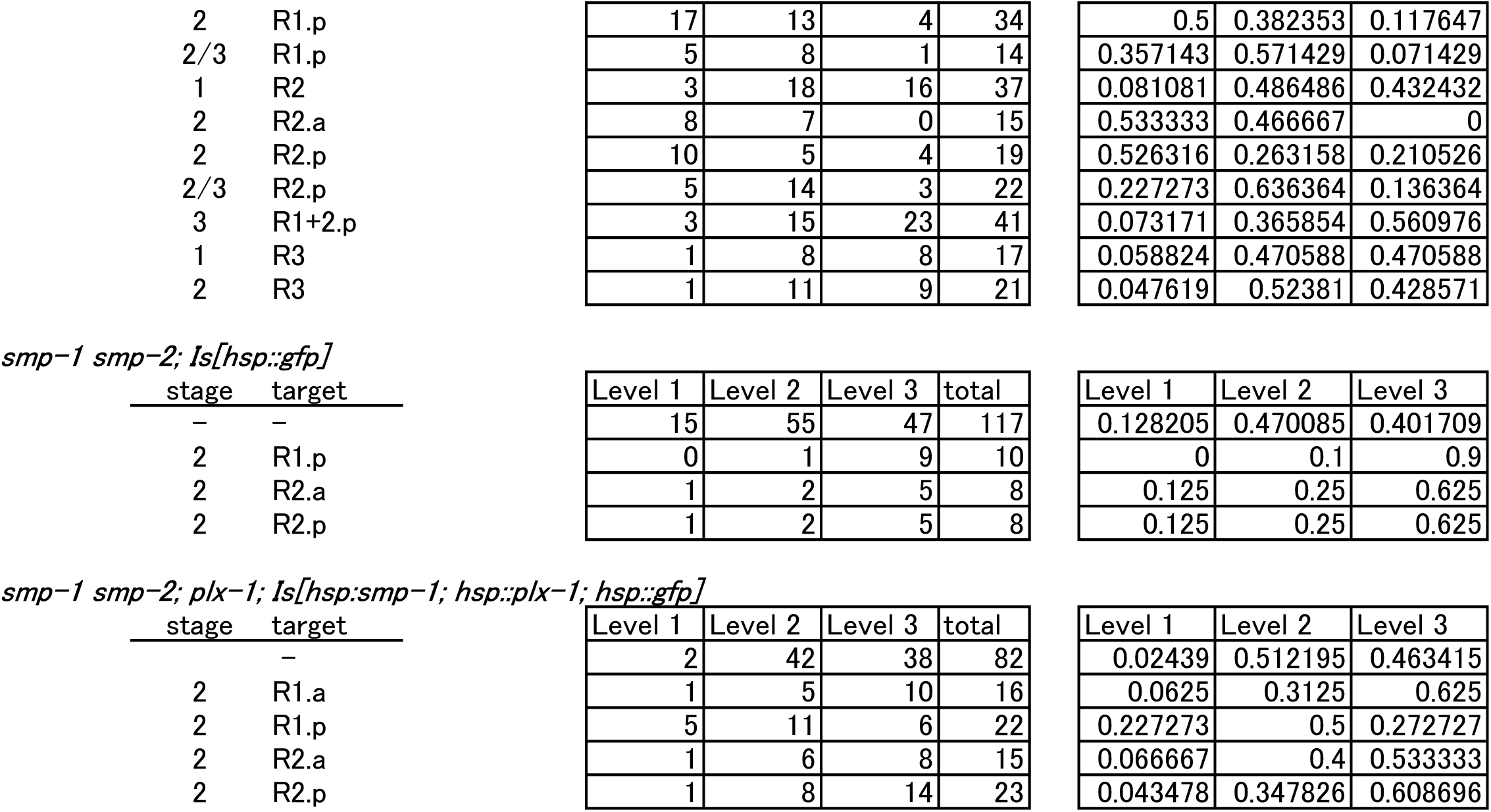
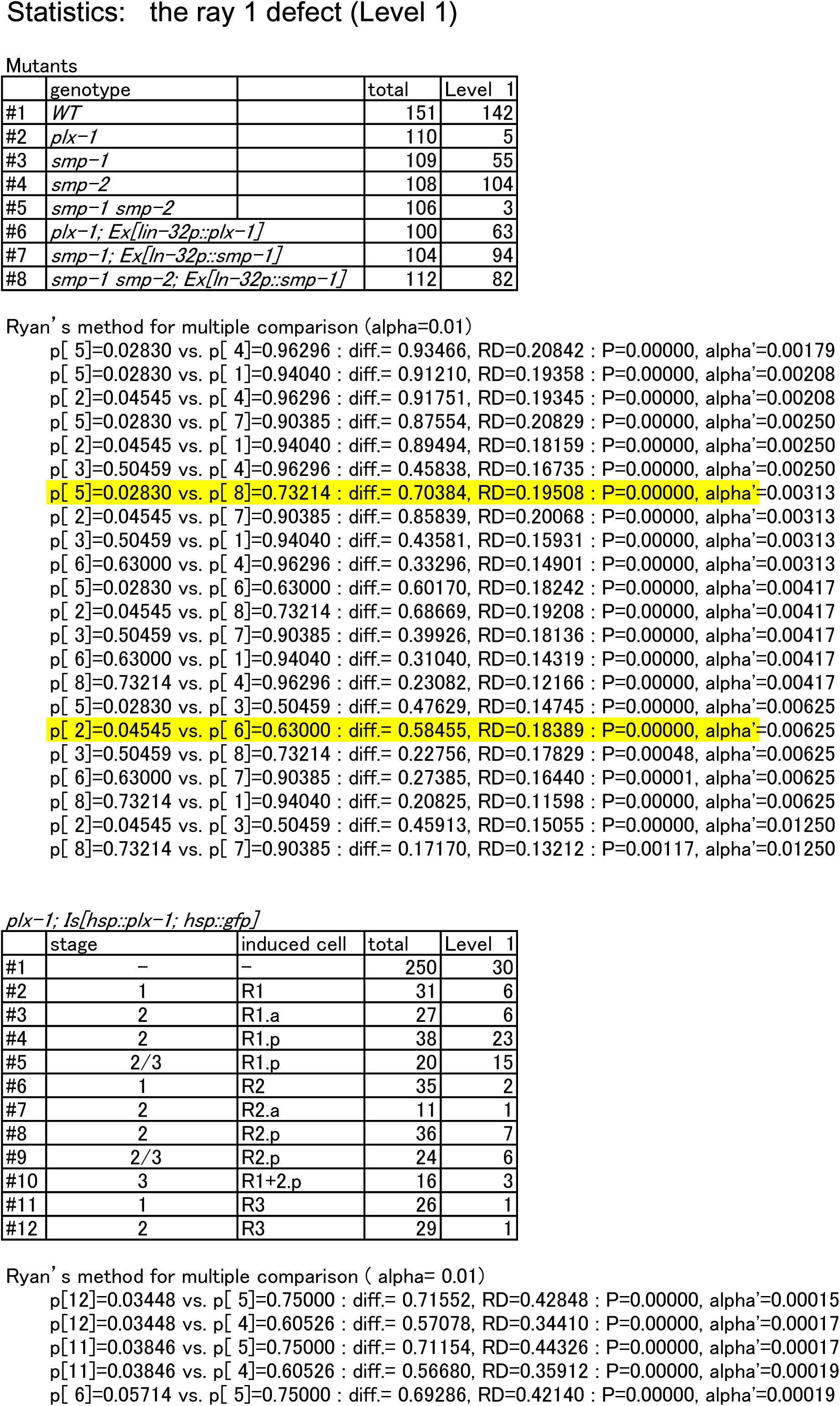
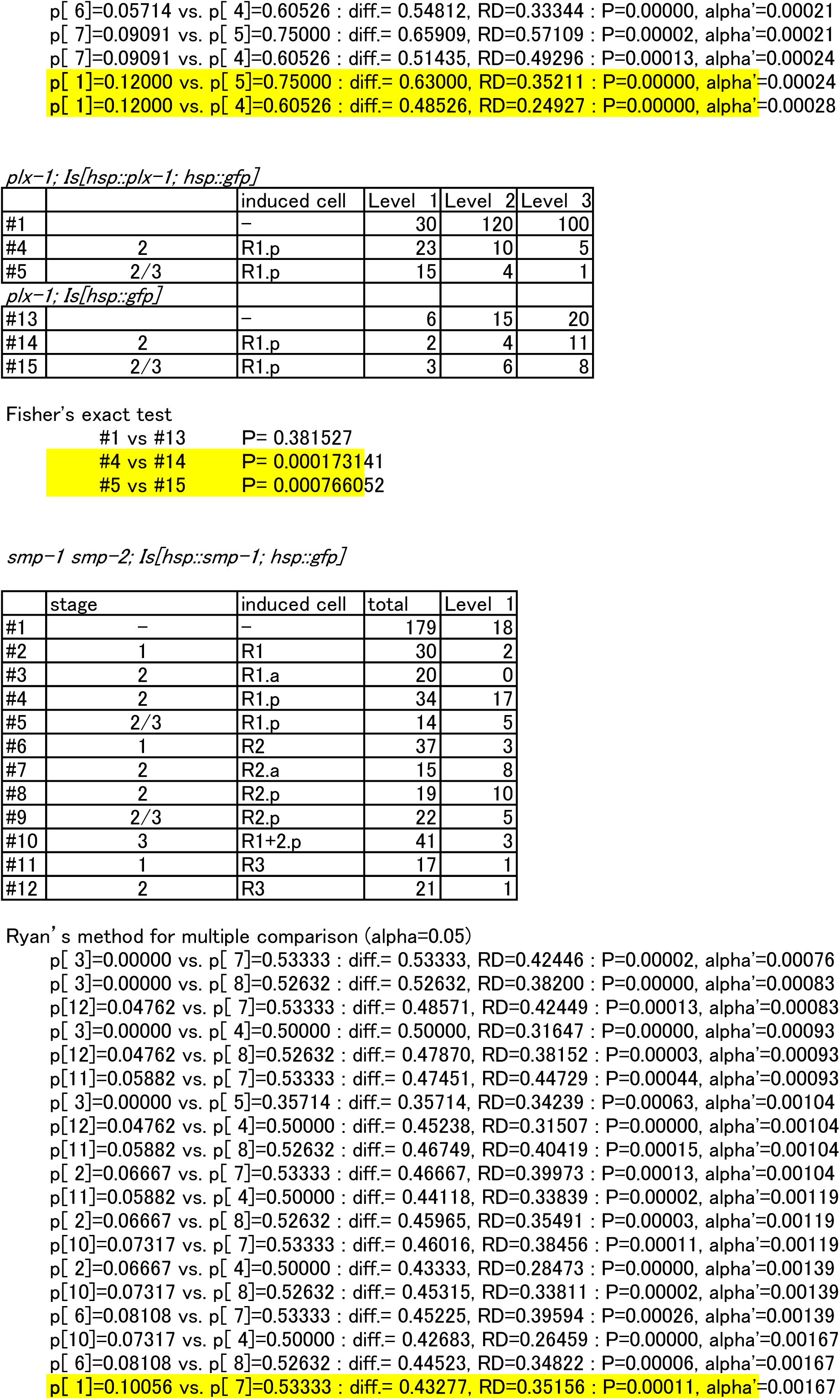
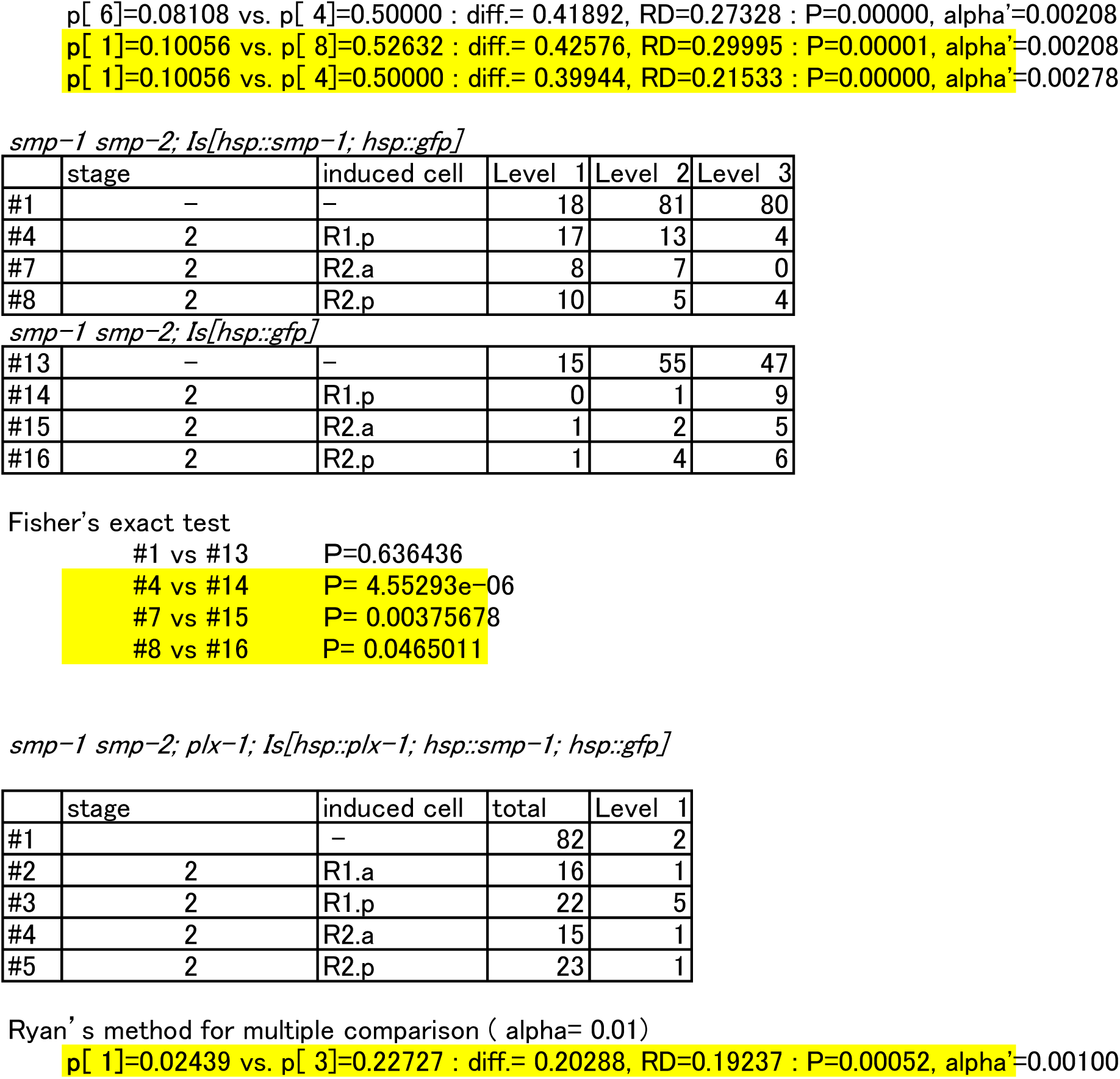
The ray 1 phenotype and the statistical analysis.

**Supplementary Table 2.** List of transgenes used in this paper.

Transgenes
|  |  |  |
| --- | --- | --- |
| *jcls1* | *ajm-1::gfp; rol-6(su1006)* | Mohler et al., 1998 |
| *ncIs13* | *ajm-1::gfp* | Liu et al., 2005 |
| *ncIs17* | *hsp::gfp* | Kamei et al., 2009 |
| *ncIs19* | *hsp::plx-1; hsp::gfp rol-6(su1006)* | Suzuki et al., 2022 |
| *ncIs23* | *hsp::smp-1; hsp::gfp rol-6(su1006)* | Suzuki et al., 2022 |
| *ncIs24* | *hsp::smp-1; hsp::gfp; rol-6(su1006)* | This paper |
| *ncIs210* | *hsp::plx-1; hsp::smp-1; hsp::gfp; rol-6(su1006)* | This paper |
| *ncIs211* | *hsp::gfp* | This paper |
| *ncEx2030* | *lin-32p::plx-1; rol-6(su1006)* | Nukazuka et al, 2008 |
| *ncEx2038* | *lin-32p::smp-1; rol-6(su1006)* | This paper |

